# Dynamic localisation of DamX regulates bacterial filamentation and division during UPEC dispersal from host cells

**DOI:** 10.1101/2021.12.17.473257

**Authors:** Bill Söderström, Daniel O. Daley, Iain G. Duggin

**Author notes:** Address correspondence to: Mail.

## Abstract

Uropathogenic *Escherichia coli* (UPEC) cells can grow into highly filamentous forms during infection of bladder epithelial cells, but this process is poorly understood. Herein we found that some UPEC filaments released from infected bladder cells in vitro grew very rapidly and by more than 100 µm before initiating division, whereas others did not survive, suggesting that filamentation is a stress response that promotes dispersal. The DamX bifunctional division protein, which is essential for UPEC filamentation, was initially non-localized but then assembled at multiple division sites in the filaments prior to division. DamX rings maintained consistent thickness during constriction and remained at the septum until after membrane fusion was completed, like in rod cell division. Our findings suggest a mechanism involving regulated dissipation of DamX, leading to division arrest and filamentation, followed by its reassembly into division rings to promote UPEC dispersal and survival during infection.

## Introduction

It is estimated that some 150 million people are affected by urinary tract infections (UTIs) each year^1^. The severity of these infections ranges from minor to life-threatening^2-4^. While there are many different species of bacteria that can cause UTIs, the major causative infectious agent is Uropathogenic *Escherichia coli* (UPEC), which is responsible for more than 80% of reported cases^5^.

The UPEC infection cycle is initiated with bacteria being endocytosed by epithelial cells in the human urinary tract^6^. Once inside the host cells, the bacteria proliferate and undergo morphological changes to adopt cocci-like shapes while forming biofilm-like Intracellular Bacterial Communities (IBC)^7^. These IBCs are exposed to various intercellular cues and develop into large communities that dominate the host cell. Eventually the colony ruptures the host cell and disperses into the surrounding environment. During dispersal, a subset of the bacteria may stop dividing and grow into long filaments, some hundreds of microns long. The filaments can later divide back into rods and reinitiate the infection cycle^8^.

It is not clear why some UPEC form filaments, but similar morphological changes have been linked to bacterial stress-responses^9-11^. It has therefore been suggested that filamentation might be a survival strategy to evade the immune system^12^. It is clear, however, that in order to reinitiate the infection cycle and colonize new host cells, the filaments must revert back to their typical rod shape^13^. As such, the reversion of filaments to rods is a crucial step in the UPEC infection cycle. To date, surprisingly little is known about how these UPEC filaments reinitiate division to revert to rods.

During vegetative growth, binary fission is mediated by the bacterial cell division machinery—the divisome^14^. This multi-protein nanomachine is organised by the highly conserved FtsZ protein^15^. Together with a number of other early arriving proteins (e.g., the essential FtsA and ZipA proteins^16,17^) FtsZ forms a proto-ring at the midcell when cells are primed to divide^18^. In a second step the divisome matures by recruiting another set of core divisome proteins that have vital functions in chromosome segregation and peptidoglycan (cell wall) synthesis ^19^. Apart from the core proteins, *E. coli* has specialised divisome-associated proteins that modify the septal peptidoglycan during division via a SPOR (*Sporulation-related repeat*) domain^20,21^. SPOR domains are widely conserved and have been found in more than 1500 bacterial genomes^22^. *E. coli*, including UPEC, has four SPOR-containing proteins^22^, DamX, DedD, FtsN and RlpA, which localise to the division septum by binding to denuded glycan strands via their SPOR domains^20,22^. Whilst the exact molecular function(s) of the SPOR proteins are yet to be elucidated, some important observations have been made. For example, DamX is involved in regulation of division of UPEC during urinary tract infections, resulting in striking morphological changes^23^. Moreover, its expression is upregulated during the dispersal (filamentation) stage of bladder cell infection and is required for UPEC filamentation, suggesting it plays a key role in arresting cell division to allow filamentous UPEC to develop^23^.

In this study, we have established an approach for studying UPEC filamentation and filament division using high-resolution and time-lapse fluorescence microscopy. And we have used this approach to quantitatively characterise the growth and division dynamics of UPEC filaments dispersed from cultured bladder cells. The data enable us to discriminate between two alternative models for the function and localization of DamX in UPEC filaments (post-infection). These models propose that DamX: (1) is localized at potential division sites along the filament during the arrest of cell division for filamentation, and then switches its function at those sites to promote division during reversal, or (2) is delocalized as part of the filamentation mechanism, and then assembles as rings, like in rod cell division, for the filament division process. The results consistently supported the second model.

## Results

### Shorter filaments are more likely to revert to rods

We used an *in vitro* urinary tract infection model system^24^, based on infection of cultured human bladder epithelial cells, to generate filaments of the model UPEC strain UTI89^25^. We transformed UTI89 with plasmid pGI5^24^, which constitutively produced cytoplasmic msfGFP and did not alter growth rate, cell size or shape in standard laboratory conditions, *i*.*e*., during growth in rich medium at 37 °C (Supplementary Figure 1). This strain was used interchangeably with wild-type (WT) UTI89 where appropriate. After infection of bladder epithelial cells with UTI89.pGI5, and then 20-22 hours exposure to a flow of urine to induce filamentation and dispersal, a sample of the flow-through containing material from ruptured bladder cells was collected, washed once in PBS, resuspended in LB, immobilized on agarose pads, and imaged with light microscopy. As observed in previous studies, there was a mixture of both filaments and rods (Fig. 1a)^13,24^ and some areas of the slide showed noticeable clusters of filaments (Fig. 1b). We defined a cell as being a filament if it was at least two and a half times longer than the average length of cells grown in liquid LB culture. Using this definition, a filament was ∼ 8 µm or more in length. The substantial heterogeneity in filament lengths (Figs. 1b and 1c) was consistent with the expected asynchronous dispersal of UPEC from the bladder cells^24^.

**Figure 1.**
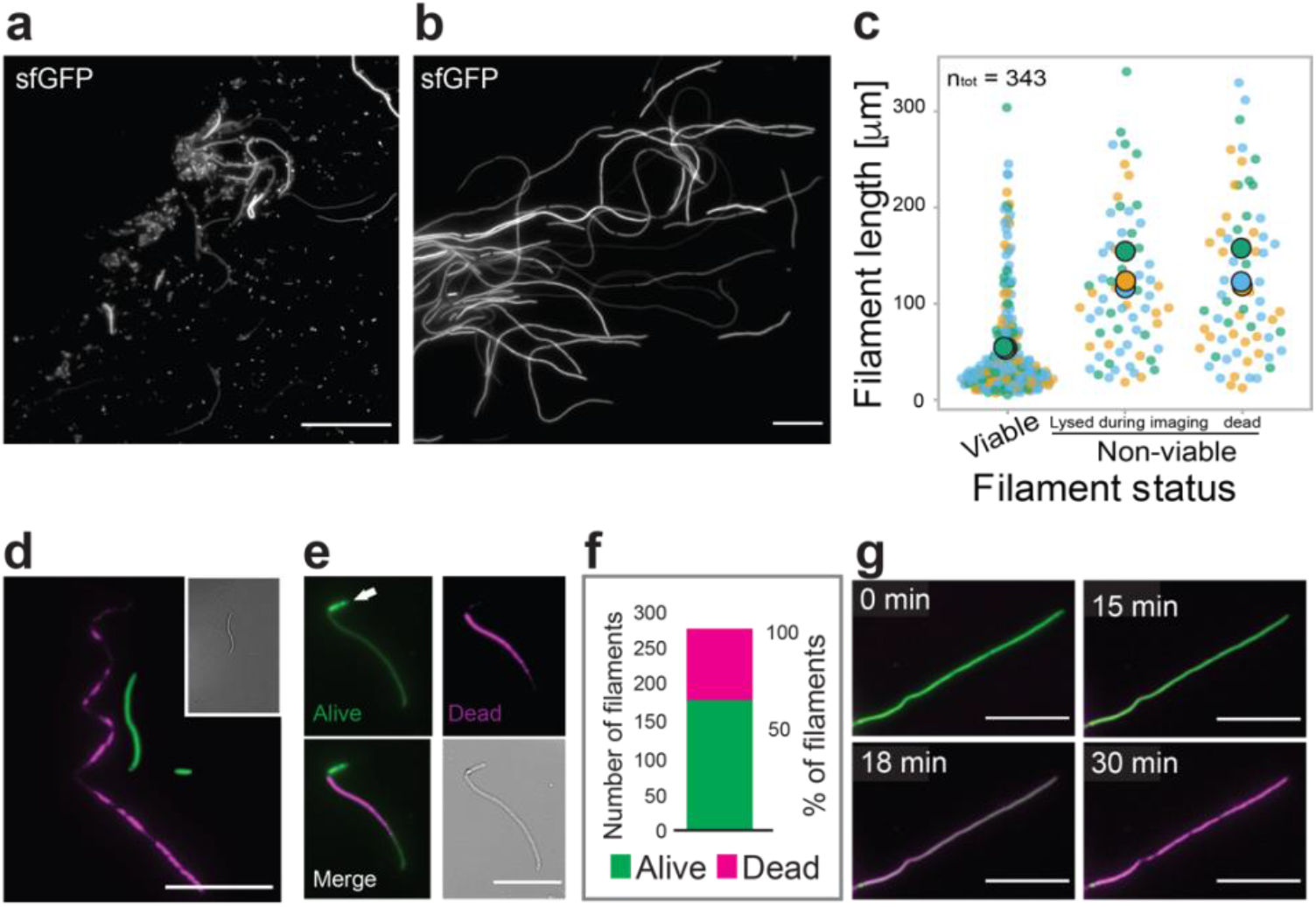
Shorter filaments are more likely to be viable and revert to rod cells. **a**, Fluorescence microscopy image showing a mixture of msfGFP-expressing short cells and filaments released from bladder cells after urine exposure. **b**, Length heterogeneity in a cluster of filaments. **c**, Shorter filaments are more likely to be viable after a round of infection. A total of 343 filaments were analysed from time-lapse imaging of three different infection experiments (Blue, Green and Yellow dots, respectively). Large, coloured dots represent the average of the respective experiment. Only cells classified as “filaments” (i.e., equal to or longer than 8 µm) were included in this analysis. **d**, WT UTI89 filaments were differentially stained to assess viability (LIVE/DEAD, green and magenta, respectively). **e**, Filaments that divided at least once during the first 120 min were classified as viable (white arrow). **f**, A total of 275 filaments from 3 infection experiments were analysed with LIVE/DEAD staining. **g**, A representative dying filament, transitioning from green to magenta over time (see Supplementary Movie SM5). Scale bars A = 40 µm, Rest = 20 µm.

To characterize the growth and division dynamics of UPEC filaments, dispersal samples were placed on a LB-agarose pad and then time-lapse microscopy movies were captured with one image acquired every 3 - 5 minutes for at least 120 min. We observed several outcomes in the filaments. Relatively shorter filaments often grew into longer filaments (> 20 µm) before starting to revert to rods, whereas some filaments elongated and then suddenly lysed (Supplementary Movie SM1). In contrast, many of the initially longer filaments (> 50 µm) did not elongate further (Supplementary Movie SM2). Most of the shorter filaments (< 50 µm) reverted to multiple rods over the course of imaging (Supplementary Movies SM3).

To quantify the state of filaments, we classified the fates of 343 filaments (from three different infection experiments) over at least 120 min into one of three groups: (1), ‘viable’, if they divided at least once, (2) ‘Lysed during imaging’, indicating filaments that did not divide and lysed during the image acquisition time (Supplementary Movie SM4), and (3) ‘dead’, representing shells of filaments that were highly translucent and did not divide nor extend in length at all during the imaging period. Supplementary Figure 2 shows examples of ‘viable’ and ‘dead’ filaments. Average lengths of filaments at the start of imaging and their classification frequencies were: viable = 56.3 ± 56.5 µm (n = 208, ∼61%), growing but no division = 117.6 ± 73.4 µm (n = 67, ∼20%) and dead = 122.6 ± 79 µm (n = 68, ∼20%) (Fig. 1c). Therefore, most of the viable filaments were significantly shorter than the average, indicating that extensive filamentation is a significant risk or stress response.

To further investigate filament viability, we used a LIVE/DEAD differential staining assay (Supplementary Figure 3) in which live cells are distinguished by their uptake of the SYTO9 green dye whereas only dead cells take up the Propidium Iodide (PI) red dye (Fig. 1d and 1e, PI shown in magenta). We examined 275 UTI89 filaments from 3 independent infections and found that 36% stained dead (Fig. 1f), similar to the proportion of non-dividing and dead filaments classified above (∼39%). We were also able to follow filaments dying over time, transitioning from green to magenta (Fig. 1g, Supplementary Movie SM5). From here on, we only characterized filaments that were classified as viable.

### Filaments divide with increasing frequency during reversion to rods

When further analysing the time-lapse movies of the UTI89 filaments, we noticed large differences in the time it took for filaments to initiate division and revert to rods, *i*.*e*., the time from placing filaments on the agarose pads until the first division event. This varied from just a few minutes to more than 2 hours in the same sample, suggesting a large heterogeneity in metabolic states between filaments. Moreover, filaments elongated by less than one micrometre, to more than 140 micrometres before the first division (mean = 19.78 ± 22.3 µm, n = 101, Fig. 2a and 2b). The time-averaged elongation rate (ΔL/Δt) varied from 0.08 µm min^-1^ (similar to standard *E. coli* growth in rich medium) to 1.72 µm min^-1^ in some filaments, with an overall mean of 0.55 ± 0.4 µm min^-1^ (Fig. 2c). We were also curious to estimate the specific growth rate (*i*.*e*., elongation rate normalized by length, Fig. 2d) and see if the elongation rate would correlate with the starting length of the filaments, however we observed poor correlation (Fig. 2e), consistent with the large heterogeneity of metabolic states of filaments (Fig. 2a and 2d).

**Figure 2.**
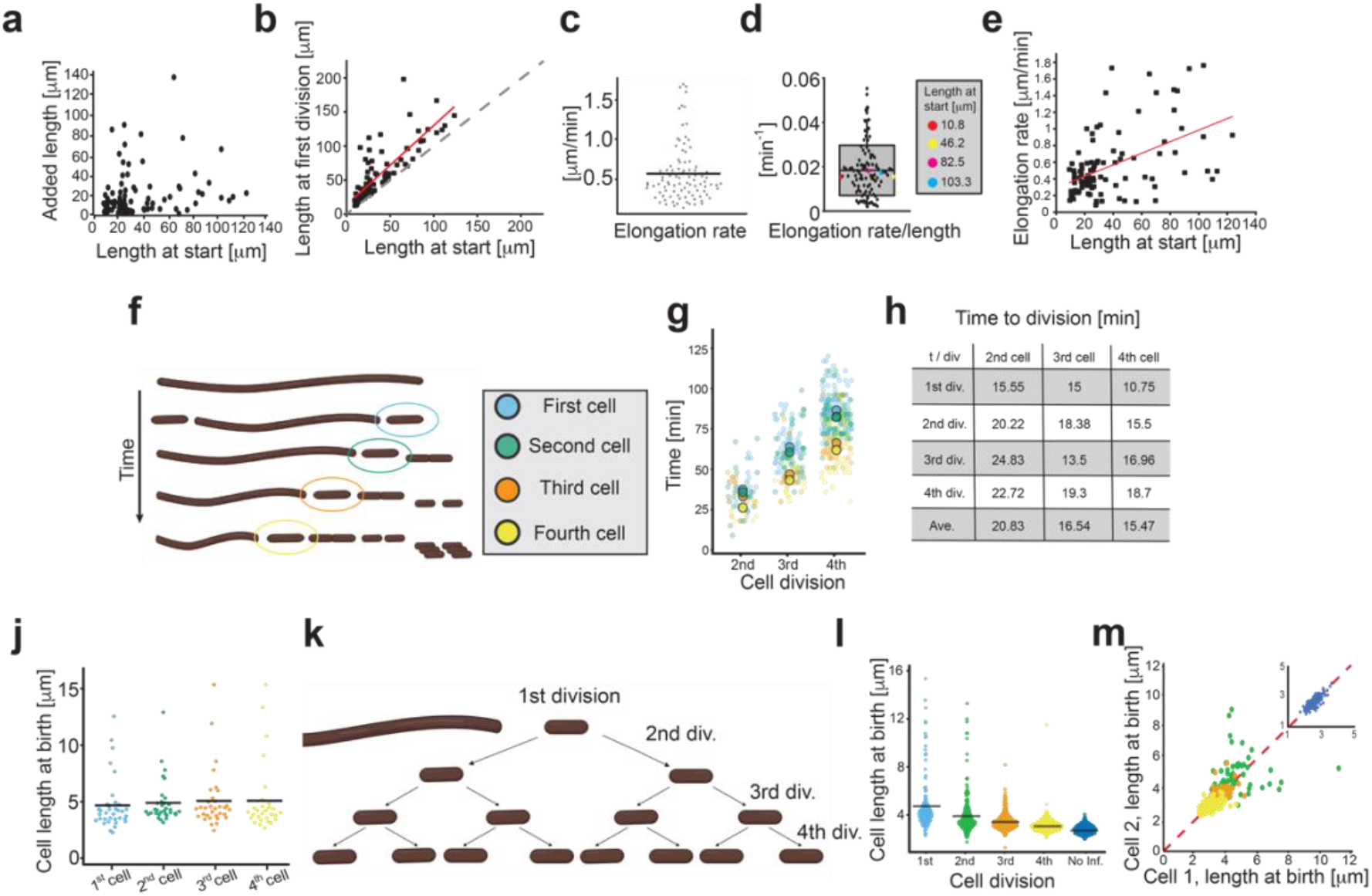
Growth and division dynamics of UTI89 filaments that revert to rods. **a**, Added cell length from the start of imaging to the first division (mean = 19.78 ± 22.3 µm, n = 101) vs cell length at the start. **b**, Relationship between filament length at start and at first division. The red line indicates a linear fit to the data. Dotted line indicates no growth before first division. **c**, Variation in elongation rate between filaments (between 0.08 and 1.76 µm min^-1^). The mean elongation rate was ΔL/Δt = 0.55 ± 0.4 µm min^-1^, n = 101. **d**, Elongation rate (0.0187 ± 0.0113 min^-1^, n = 101) of filaments normalized to their length at start of the imaging. Box indicates SD, midline indicates Mean, whiskers indicate 95% interval. Box to the right high-lights four randomly picked filaments with rates close to mean (red = 0.015 min^-1^, yellow = 0.0152 min^-1^, magenta = 0.0178 min^-1^, blue = 0.017 min^-1^), these filaments ranged from 10 to 100 um in length, indicating weak correlation between length and elongation rate. **e**, The elongation rate was not correlated to the length of the filaments at the start of imaging. The red line shows a linear fit to the data. R^2^ = 0.245. **f**, Schematic representation of the first to fourth generations of cells during filament division. **g**, Timing of subsequent division events in ‘mother’ filaments after birth of the newborn cells depicted in panel F (colour key). Larger circles represent the means (n_div_ = 471). **h**, Mean inter-division times in subsequent divisions of the mother filament after the 2^nd^ 3^rd^ and 4^th^ cells have pinched off. Overall means were: 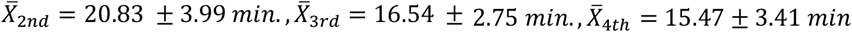. (average ± S.D.) **j**, Mean cell lengths of the smaller newborn cell pinched off from a filament (n = 135). **k**, Schematic representation of the subsequent division events of pinched-off newborns. **l**, Lengths at birth of daughter cells from first to fourth generations. **m**, Symmetry of division in newborns. Second generation (green) cells divide more asymmetrically than third (orange) and fourth (yellow) generation of cells. Inset; UTI89 grown in LB divides highly symmetrically. All values represent Mean ± SD.

We next monitored individual filament division over four consecutive generations, after which most cells became too crowded and moved out of focus (Supplementary Movie SM6). Most filaments grew initially (Fig. 2b) and then often divided towards one end, resulting in rod cells ‘pinching-off’ (Fig. 2f and Supplementary Movie SM6). The sequential generations of the ‘mother’ filament occurred at progressively greater frequency (Fig. 2g), often appearing at multiple locations near both ends. This was also be seen by measuring the mean filament interdivision time (Fig. 2h), where, for example, there was a ∼25 % difference between filament division frequencies after the second and the fourth cells had pinched off (second cell, 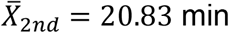, compared to the fourth cell, 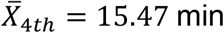). The short interdivision times would suggest that multiple potential division sites may be primed as soon as—or even before—the previous generation has pinched off from the mother filament.

The size of newborn cells pinching off from the filament were quite consistent over time, with mean lengths of 4.66 ± 2.32 µm for the first cell, and then 4.89 ± 1.92 µm, 5.05 ± 2.59 µm, and 5.08 ± 3.1 µm for subsequent generations. There was a noticeable size skew, however, seen at all generations, with ∼80% of newborns shorter than average and ∼10% of newborns at least twice the average length (Fig. 2j and Supplementary Movie SM6). We then tracked the size of each pinched-off cell for 3 - 4 of its subsequent divisions (see Fig. 2k), which revealed a slight decrease in mean length over four generations: 4.83 ± 2.33 µm (n = 129), 4.01 ± 1.71 µm (n = 203), 3.44 ± 0.67 µm (n = 397) and 3.11 ± 0.5 µm (n = 664), respectively (Fig. 2l). For comparison, cells that had not been through an infection cycle were on average 2.79 ± 0.35 µm (n = 340) in length at birth (Fig. 2l, ‘No inf.’). The first and second divisions sometimes gave rise to abnormally long cells (Fig. 2l), but this corrected over the next one or two generations, with essentially all pairs of daughter cells in the fourth generation being the same length at birth (Fig. 2m).

### Nucleoid distribution during filament reversal

To see how DNA partitioning was maintained during reversal, we also looked at the localization of chromosomes in filaments of UTI89 transformed with a plasmid expressing HupA-RFP^26^ (which did not interfere with UTI89 growth; Supplementary Figure 1). Fluorescence microscopy revealed that the DNA was distributed along the lengths of filaments and in somewhat irregular patches (Supplementary Figure 4A, 0 min). Filaments contained on average 28.2 nucleoids per 100 µm cell length, corresponding to ∼3.5 µm cell length per nucleoid, which is similar to normal rods. The average length of nucleoid masses in filaments without visible constrictions was 2.05 ± 0.86 µm (n = 305), with a primarily bimodal size distribution (Supplementary Figure 4B, red). In post-divisional rods that had been pinched off from a filament, and were about to divide, the average nucleoid length was 1.3 ± 0.24 µm (n = 254) (Supplementary Figure 4B, green), similar to non-infection UTI89 growing in LB of 1.45 ± 0.32 µm (n = 308) (Supplementary Figure 4B, blue). There was an increasing degree of symmetry in chromosome partitioning over generations (Supplementary Figure 4B, Inset), consistent with the observed progression of cell size symmetry (Fig. 2k).

### Localization of DamX in filaments reverting to rods

Our results above showed that filaments surviving the infection conditions rapidly coordinate reversion to rods when conditions become favourable for vegetative growth. It is not known how or when the cell division machinery assembles in UPEC filaments to reinitiate division. The non-essential *E. coli* division protein, DamX, is required for infection-induced filamentation of UTI89^23^. DamX is localized to the divisome during vegetative growth in an strictly FtsZ-dependent manner^27^. To visualize DamX in filaments and during their subsequent reversal, we produced a fluorescent protein fusion, mEos3.2-DamX, expressed from a plasmid, in UTI89. In our experimental conditions, the fusion represented ∼60% of the total cellular DamX (Supplementary Figure 1C), which did not affect growth (Supplementary Figure 1A-B), consistent with previous studies on over-expression of cell division proteins ^28-30^. Furthermore, localization in rods after infections were similar to what has been seen in K-12 strains^22^ (Supplementary movie SM7). High-level DamX overexpression has previously been shown to cause filamentation^31,32^, and its strong upregulation is associated with the UTI89 filamentous response during infection. We found that the moderate production of mEos3.2-DamX did not inhibit filament reversal.

Filaments obtained from infections were imaged using live cell single-molecule PhotoActivated Localization Microscopy (PALM) and epi-fluorescence time-lapse microscopy. The resulting images indicated that mEos3.2-DamX localized as broad rings prior to visible invagination (Fig. 3a and 3b). Indeed, measurement of the diameter of the mEos3.2-DamX rings and of the filament 1 µm either side showed a ratio of 1:1.05 – 0.95 (Fig. 3c and 3d). The fluorescence intensity profiles along the filament axis showed that the pre-constriction mEos3.2-DamX localizations were as wide as > 500 nm (Fig. 3c and 3e). During division, the diameter of the mEos3.2-DamX rings naturally narrowed to smaller radial width than the membranes (Fig. 3f and 3g). The mean axial breadth of condensed mEos3.2-DamX rings along the cell length was 116.5 ± 13.4 nm (n = 122) (Fig. 3h), similar to those of cell not run through an infection (102.5 ± 20.2 nm, n = 150) (Supplementary Figure 5). Our data also indicated that, similar to another SPOR-domain protein, FtsN^33^, mEos3.2-DamX seemed to linger at the division sites after the inner membranes had closed, before being completely disassembled (Fig. 3j-l).

**Figure 3.**
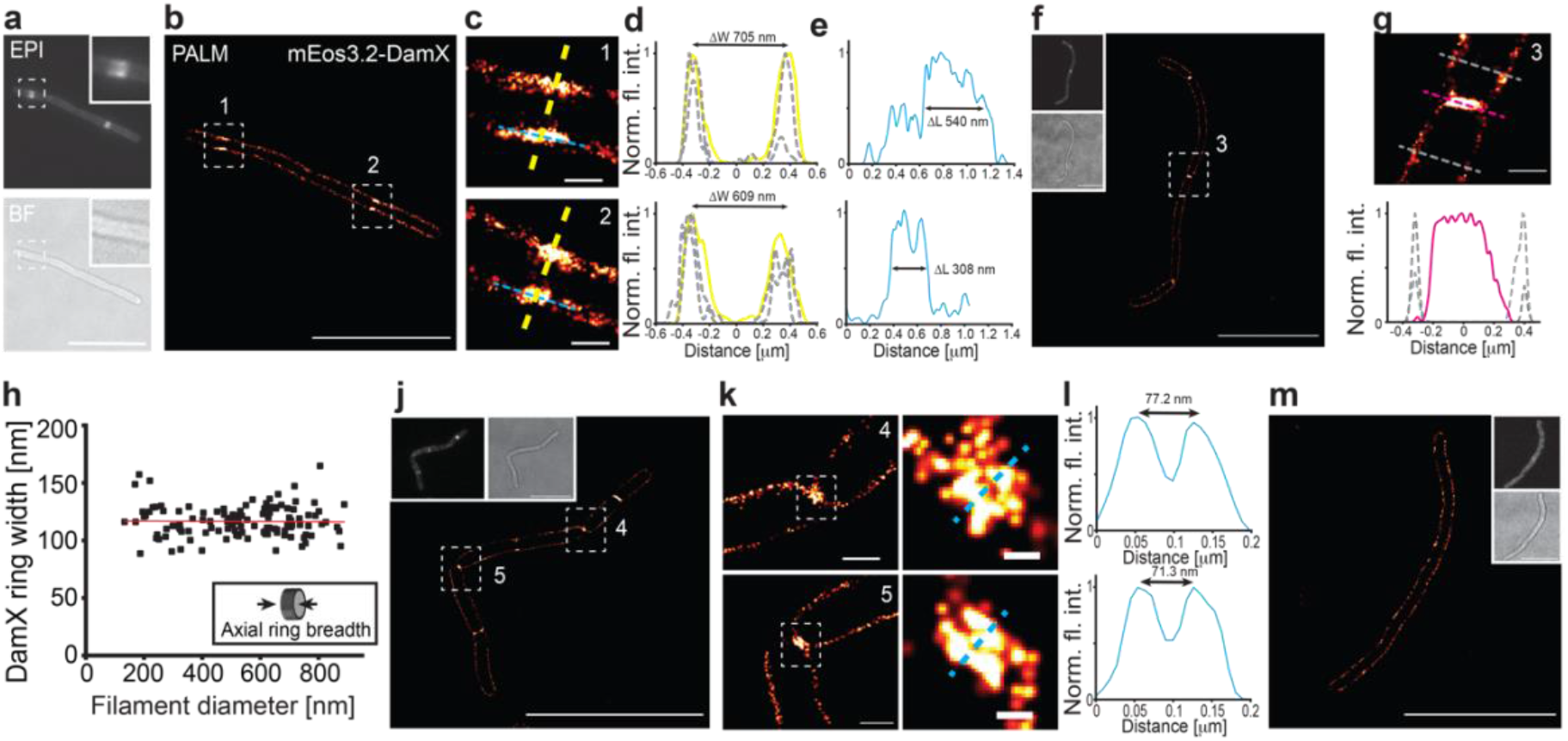
DamX localises at the division site prior to membrane constriction. Filaments expressing mEos3.2-DamX harvested from the back-end of flow chambers after infections were grown in LB for 1 hour before imaged using single-molecule microscopy. **a**, mEos3.2-DamX localized at multiple division sites simultaneously and accumulated prior to visible invagination of the membranes. Insets: Close-up images of mEos3.2-DamX and the corresponding brightfield image. **b**, PhotoActivated Localisation Microscopy (PALM) image of the same filament as in **a. c**, Close up images (1) and (2) of the mEos3.2-DamX ring assemblies at division sites prior to condensing into a ring structure (from panel **b**). **d**, Fluorescence intensity plots of the mEos3.2-DamX assemblies and membrane widths. Plots showing ring and membrane widths; yellow lines represent ring assemblies; grey dotted lines represent membrane widths 1 µm up and downstream of the mEos3.2-DamX accumulation. ΔW indicates the peak-to-peak distances. **e**, Length of the molecule assemblies along the length axis of images in **c**. ΔL indicates the length of the intensity profile at 50% of the intensity. **f**, PALM image of a typical filament during reversal. **g**, Close up image of a constricting mEos3.2-DamX ring (3), fluorescence profile underneath: yellow line represents the ring, grey represents membrane width. **h**, Axial breadth of mEso3.2-DamX rings along the length of the filaments at various cell diameters. Average width 116.5 ± 13.4 (n = 122), values represent Mean ± SD. Red line represents linear fit to the data: y = −0.009 ∗ x + 116. **j**, mEos3.2-DamX remains at the old division septum after membrane separation. **k**, Close ups of old division sites from panel **j** (4) and (5). **l**, Peak-to-peak distance of fluorescence intensities of the membrane assemblies of mEos3.2-DamX. **m**, A typical filament before division sites have been defined. Scale bars (**a, b, f, j, m**) = 10 µm, (**c, g, k**_**left**_) = 500 nm, (**k**_**right**_) = 100 nm.

Filaments obtained from infections that were alive but not yet primed for division did not have mEos3.2-DamX rings (Fig. 3m). We then used time-lapse epifluorescence imaging, which showed that mEos3.2-DamX assembled as rings at different places along the length of the filament during the image acquisition period (Supplementary movie SM8). Figure 4A shows the formation of at least three generations of division rings in one filament over time, and importantly, that the filaments then reverted into rods at those sites. Once an mEos3.2-DamX ring formed, it remained in place until division was completed. Measurement of mEos3.2-DamX ring position indicated that there was a preference for rings to form initially at the approximate ¼, ½ or ‘positions along the filament length regardless of the length, which was predominantly in the range of 7.5 – 40 µm (n = 134) (Fig. 4b). The data also indicated that cells pinching off in the second and third generation did so closer to the poles (Fig. 4b and 4c). Note that the mother filament would have grown in length between each division.

**Figure 4.**
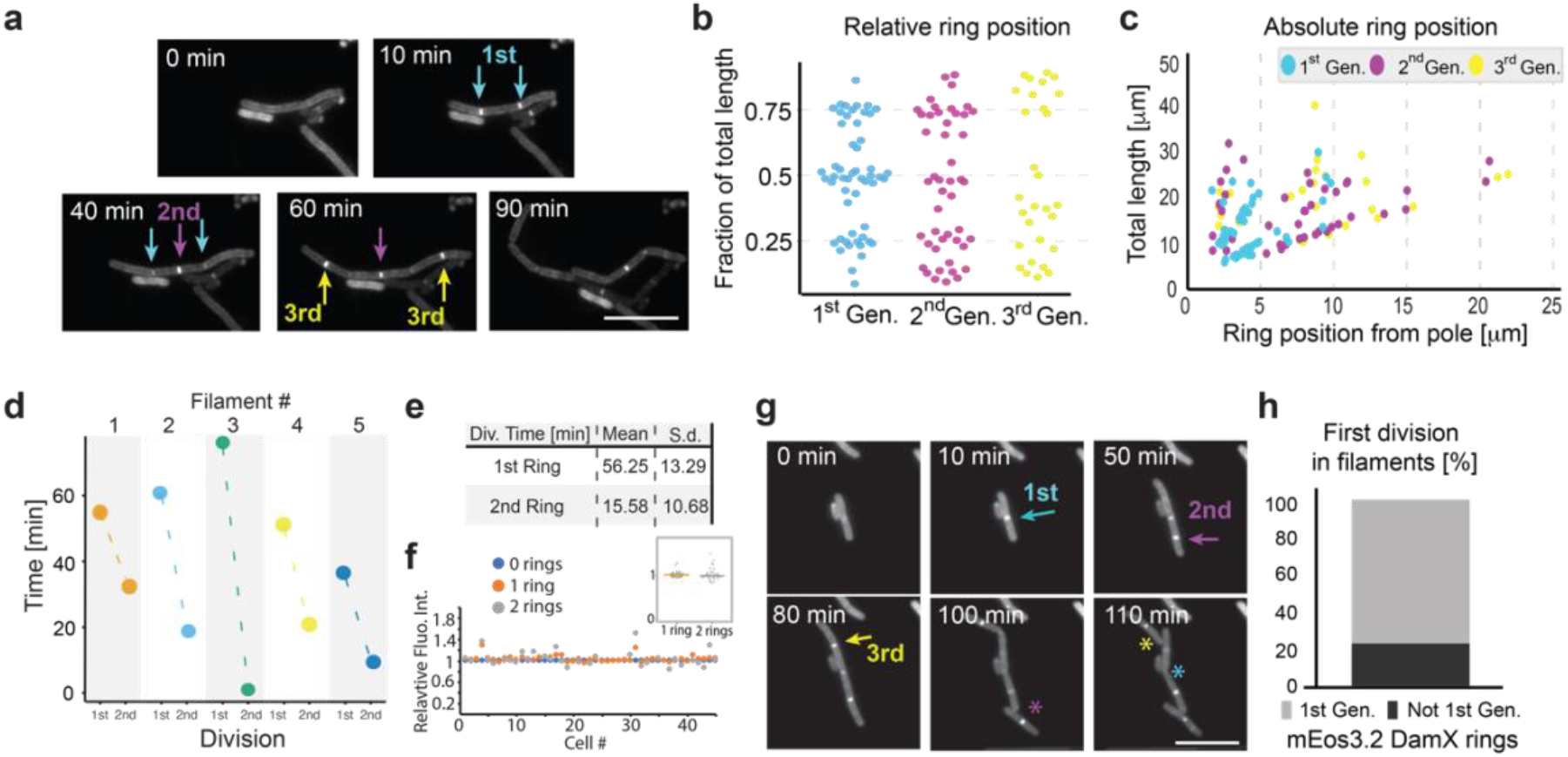
mEos3.2-DamX localization and dynamics in filaments. **a**, Time-lapse images of filaments expressing mEos3.2-DamX (Supplementary movie SM9). Formation of mEos3.2-DamX rings were followed over time in filaments where no rings were observed in the first image. Cyan arrows show 1^st^ generation, magenta arrow 2^nd^ generation and finally yellow arrows indicate the 3^rd^ generation division rings. **b**, Relative de novo positioning of mEos3.2-DamX division rings in filaments. Most rings assembled at locations close to 1/4, 1/2 or 3/4 of the total filament length. Colour coding follows that in panel **a**. Gen. = Generation. n = 134. **c**, The distance of mEos3.2-DamX rings in µm from one of the cell poles. **d**, Plot shows the time from first mEos3.2-DamX ring formation to the first division (1st), and the time from the first division to the second (2nd, Δt = t_2_ - t_1_) for five randomly picked filaments. **e**, Summary showing average times of first and second division based on mEos3.2-DamX ring formation and constriction. **f**, Total cellular mEos3.2-DamX fluorescence intensity did not change with formation of division rings. Blue dots represent total integrated fluorescence normalized to filament area one frame before formation of the first ring, orange dots represent integrated fluorescence when one ring had formed, grey dots represent integrated fluorescence when two rings had formed. **f inset**, Relative mean cellular fluorescence with one ring = 1.02 ± 0.07 (n = 45), relative mean cellular fluorescence with two rings = 1.03 ± 0.15 (n = 32). Values represent Mean ± SD. **g**, A shorter filament from a time-lapse movie indicating that the second generation of mEos3.2-DamX ring (magenta, formed at t = 50) constricted and pinched off prior to the first ring formed (cyan, formed at t = 10 min). First image showing a full division is at t = 100 min, at the place where the second mEos3.2-DamX ring was formed (indicated by a magenta asterisk). Division of the first mEos3.2-DamX ring is indicated by cyan asterisk. A third-generation mEos3.2-DamX ring is indicated by the yellow arrow (with division indicated by yellow asterisk). Scale bar 10 µm. **h**, 23 % of the second generation mEos3.2-DamX rings formed in filaments pinched off before the first generation mEos3.2-DamX rings. All scale bars = 10 µm.

In line with our observations that the apparent time between division of daughter cells from filaments decreased with successive generations, we wanted to know if this was also consistent in the first division (previously, we could only determine the relative time from the second division without a division marker, *i*.*e*., Fig. 2). That is, would the first division ring be present longer than the second before a division? To do this, we followed mEos3.2-DamX ring formation, constriction, and subsequent division in filaments over time. Indeed, the average time from the first mEos3.2-DamX ring formation to the first division was 56 ± 13 min (SD), and there was a significant reduction in time to the next (second) division: 15 ± 10 min (SD) (Fig. 3d-e).

We also wanted to see if the intensity of mEos3.2-DamX varied during ring formation, as this would be proportional to the total cellular expression level. The total cellular fluorescence did not change significantly with formation of new mEos3.2-DamX rings (Fig. 4f); compared to filaments with no visible rings, the relative mean cellular fluorescence with one ring was 1.02 ± 0.07 (n = 45), while relative mean cellular fluorescence with two rings was 1.03 ± 0.15 (n = 32) (Fig. 4f, inset). Curiously, we also noticed that the mEos3.2-DamX rings did not always divide in the order that they initially formed (Figure 4G). It was observed that around 20% of filaments had the second generation of rings pinch off prior to the first (Fig. 4h).

Since mEos3.2-DamX is assembled at multiple sites along filaments, we wondered whether septal PG (sPG^34^) synthesis also occurs at multiple sites at the same time. To test this, we pulse-labelled filaments during reversal to detect regions of sPG synthesis with an OregonGreen488-labelled Fluorescent D-amino acid (OGDA)^35,36^. As we saw with mEos3.2-DamX, OGDA was observed in multiple fluorescent rings (Fig. 5a), indicating that active peptidoglycan synthesis was present at multiple sites and that multiple cell division machineries were active in multiple rings simultaneously.

**Figure 5.**
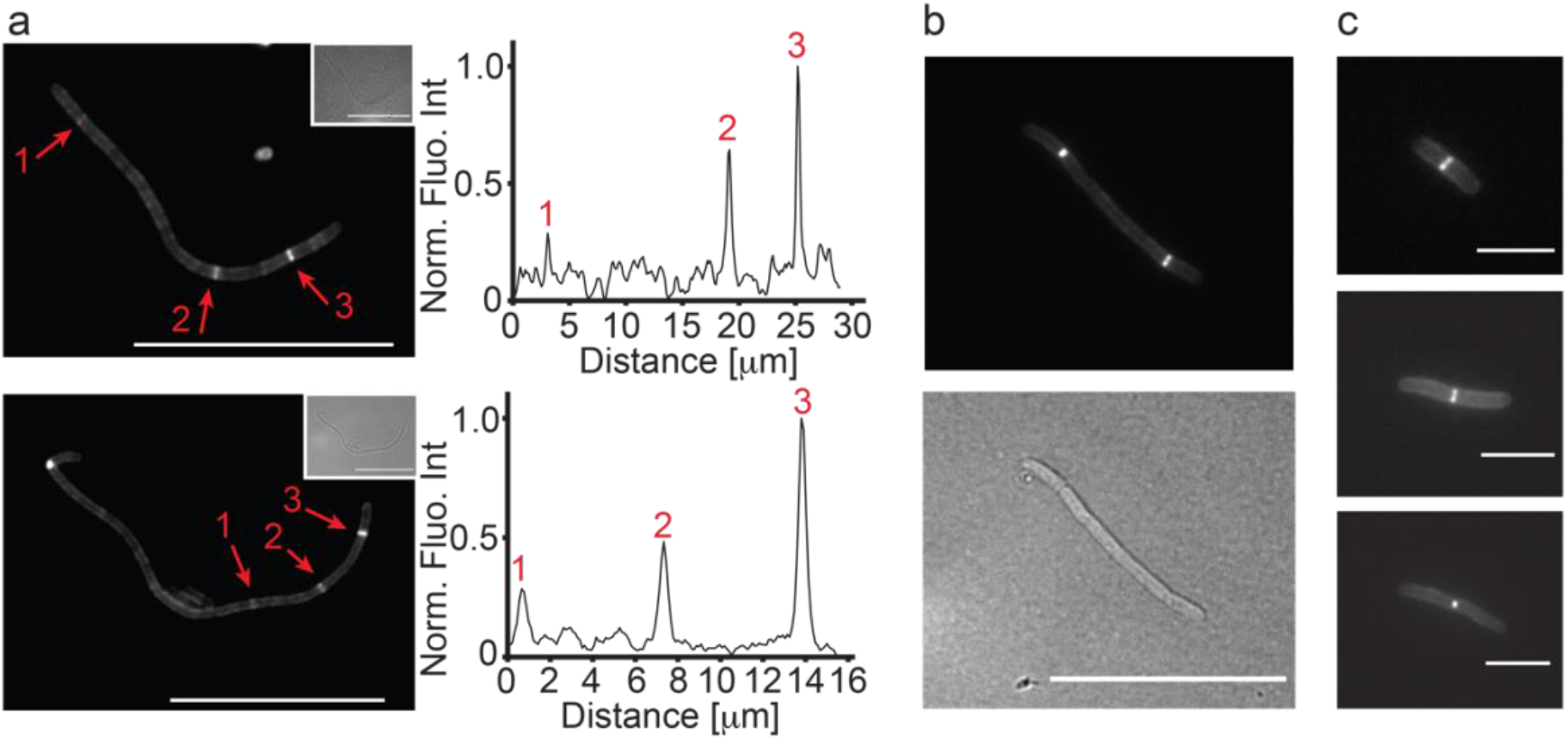
Peptidoglycan synthesis is active at multiple locations at the same time. WT UTI89 filaments from an infection cycle were labelled with the green fluorescent D-amino acid (FDAA) probe OGDA and imaged using fluorescence microscopy. Harvested filaments were pelleted and resuspended in LB and placed at 37 °C for two hours before OGDA labelling for five min. **a**, Multiple active sites for peptidoglycan synthesis can be observed in filaments. Inset: corresponding brightfield images. Fluorescence intensity plots for selected sections are shown next to the images (peaks numbered). **b**, A representative mid-length filament with two clear OGDA accumulations at division sites. **c**, Rod shaped cells in various stages of membrane constriction. Cells had already reverted from filaments and were then stained with OGDA, showing fluorescent staining only at midcell, as expected. Scale bars **a** and **b** = 20 µm, **c** = 4 µm.

We also observed various levels of OGDA intensity, indicating differing activity of sPG synthesis in the rings. The localisation of OGDA at many division sites preceded visible constriction, suggesting that the cell division machineries would be assembled at that stage, whereas other OGDA localizations were clearly associated with later-stage division constrictions in both filaments and rods (Fig. 5b and 5c).

## Discussion

We have used live cell time-lapse, epifluorescence and super-resolution microscopy to quantitatively characterize the growth and division of filamentous Uropathogenic *E. coli* (UPEC) released from the dispersal stage of infection of bladder epithelial cells^24 25^. UPEC filaments can grow to hundreds of microns long in UTI, but we found that shorter filaments (< 50 µm) were more likely to survive and revert to rods in culture. UPEC filamentation has been proposed as a survival strategy to prevent their potential phagocytosis by immune cells or improve surface adhesion^10,11,13^. Our results suggest that UPEC filamentation is a response to stress during dispersal and exposure to human urine—and the stress is not always overcome.

Prior to filament reversal (i.e., division into rods) most filaments undertook a period of variable growth (on LB agarose), where elongation rates ranged from 0.08 µm min^-1^ to 1.72 µm min^-1^, or up to ∼20 times the elongation rate of a standard rod cell. Since filamentation combines the growth power of multiple rods and effectively links the strength of their cell walls together in one cell body, we propose that the rapid directional elongation of filaments physically aids UPEC dispersal from the IBC biofilm and host cell carcass, helping them to ‘push out’ into the extracellular environment. Indeed, the extrusion-like images of UPEC filaments exiting host cells are consistent with this potential selective advantage^24,37^.

The elongation rate of UPEC filaments was not correlated with the initial length of the filament, indicating a substantial variability in metabolic states of filaments regardless of length. Consistent with this, the time to the first division ranged from just a few minutes to more than two hours. The substantial heterogeneity may be related to both the asynchronous dispersal from the bladder cells and the different morphological fates of individual IBC dispersal events^11,13,24^.

Most UPEC filaments divided near the ends, releasing a smaller ‘pinched-off’ cell. Over the first few divisions, the filament interdivision time became shorter, much like filament reversal after antibiotic treatment^38^. Furthermore, the pinched-off UPEC cells became shorter over the initial generations too, approaching the size of standard rods. The combined results suggest that division tends towards balanced growth over a few generations. Previous studies looking at reversal of *E. coli* filaments induced by antibiotics and DNA damage (SOS response) found that division rings assemble evenly along the cell, sometimes only transiently and then assembling at new locations, but then one division event occurred at any given time^38^. In contrast, we found that in infection-induced UPEC filaments, multiple overlapping divisions were common, predominantly towards the poles. This suggests that significantly different mechanisms of filamentation and reversal underpin the antibiotic, SOS-response and infection/dispersal models of filamentation and highlights the importance of understanding these differences in future development of infection therapeutics.

Reversal is a critical event for UPEC because only rods are thought to be endocytosed by host cells to initiate a new round of infection7, amplifying an acute UTI. However, the regulation and mechanisms initiating reversal are unknown. A recent study found that DamX—previously implicated in sPG remodelling during normal division—is needed for UPEC filamentation and was upregulated during dispersal^23^. This led to the hypothesis that DamX switches functions to block division at potential division sites during UPEC filamentation. Surprisingly, we instead observed that fluorescently-tagged DamX (mEos3.2-DamX) was not initially assembled at potential division sites in UPEC filaments, but assembled later as rings where filaments eventually divided.

Our single-molecule data suggest that mEos3.2-DamX assembles into rather broad rings at the quarter or half positions along filaments, consistent with the expected locations of division, and then condenses leading up to constriction, where mEos3.1-DamX rings showed very similar dimensions as those formed by related cell division components during *E. coli* division ^28,39,40^. Lastly, mEos3.2-DamX remained assembled at the division septum until after the envelopes had visibly separated, suggesting a role for DamX in sPG regulation until the new cell wall is completed. In summary, the current data suggest that during filamentation DamX has a role in blocking division that involves regulated delocalization, possibly associated with sequestration of division proteins, but then switches modes for reversal to assist in cell division in a manner that structurally resembles normal cell division.

## Methods

### Bacterial cell growth

A single colony of respective *E. coli* UTI89 strain was grown overnight in a 20 ml culture at 37 °C without shaking, to favour expression of type-1 pili which facilitates adhesion to the bladder cells during infection^5^. Antibiotics were added when needed (ampicillin 100 µg ml^-1^, spectinomycin 25 µg ml^-1^). The following morning, the cultures were pelleted and resuspended in PBS to a concentration of OD_600_ 0.2 and added to the infection model. Cells not used in the infection model were diluted 1:50 and grown to an OD_600_ of ∼ 0.4 before imaging to measure cell length and fluorescence profiles.

### Plasmid construction

The expression plasmid pAJM.011 was used as a backbone in the construction of pDD7 (mEos3.2-DamX). This plasmid was part of the Marionette sensor collection^41^, which was obtained from Addgene (Kit #1000000137). Initially the *aph* coding sequence in pAJM.011 was replaced by the *bla* coding sequence. In a second step, the coding sequence for EYFP was replaced by the coding sequences for mEos3.2 and DamX. All fragments were amplified by polymerase chain reaction (PCR) using Q5 DNA polymerase (New England Biolabs, USA). Fragments to be ligated contained regions of 20-30 bp of homology, and were cloned using the *in vivo* DNA assembly method^42^. The coding sequence for bla was obtained from pET15b, mEos3.2 from Genscript (The Netherlands) and DamX from the *E. coli* strain MG1655. Oligonucleotide synthesis and DNA sequencing was performed by Eurofins Genomics (Germany).

### Fluorescent protein production

WT UTI89 strain was transformed with pGI5^24^ or pDD7 to produce strains expressing msfGFP or mEos3.2-DamX, respectively. msfGFP did not need inducer as it was produced from a constitutive promoter^24^. mEos3.2-DamX expression was induced by adding 1 µM anhydrotetracycline (aTc) for 1 hour to the infection cultures prior to collecting filaments. WT UTI89 strain was transformed with pSTC011^26^ to produce a strain expressing HupA-RFP. HupA-RFP was induced for 1 hour by adding 20 µM isopropyl β-D-1-thiogalactopyranoside (IPTG) to the cultures containing filaments.

### Preparation of human urine samples

This study has human research ethics approval form the UTS Human Research Ethics Committee (HRCH REF No. 2014000452). Urine from two different donors (one female and one male) was collected in the morning and stored for at least 2 days at 4 °C. Samples were centrifuged at 4500 x g for 8 minutes and the supernatant was filtered through a 0.2 µm membrane filter. The specific gravity was determined by comparison to pure water using a gravity meter. Only urine samples in the pH range 5.12-5.83 and USG range 1.024 – 1.031 g ml^-1^ was used^24^. Filter sterilized samples were frozen at -20 °C until required. Note, to be as consistent as possible, only one batch of urine was used per technical replicates, since both pH and USG can alter the proportion of filamentation in the samples^13^.

### Urinary tract infection model

We based our infection model on a previously described approach^13,24^, with minor modifications. In summary, on day one, flow chambers (IBIDI µ-Slides I^0.2^ Luer, Cat#: 80166) were seeded with PD07i epithelial bladder cells at a concentration of ∼ 3×10^6^ cells in EpiLife Medium (Gibco) supplemented with growth supplements and antibiotics (HKGS and 100µg ml^-1^ Pen/Strep). Chambers were left overnight for cells to adhere and grow into a confluent layer. The next day, flow chambers were connected to New Era pumps via tubing and 20 ml disposable syringes, and a flow (15 µl min^-1^) of fresh EpiLife (supplemented with HKGS) without antibiotics was maintained for 18-20 hours. On day three, to induce infection, bladder cells were exposed to bacterial cultures at a concentration of OD_600_ 0.2 for 20 minutes at a flow rate of 15 µl min^-1^. Following this step, the media was changed back to EpiLife (supplemented with HKGS), after an initial flow of 100 µl min^-1^ to flush out the excess bacteria, and flowed for 9 hours to allow bacteria to adhere to and invade the epithelial bladder cells. This step was followed by flow (15 µl min^-1^) of EpiLife (supplemented with HKGS) in the presence of 100µg/ml gentamycin for 20 hours to allow for the formation of intracellular bacterial communities (IBCs), as well as to kill and wash away any lingering extracellular bacteria. Following these 20 hours, the media was changed to human urine (with pH between 5.12-5.83 and Urine Specific Gravity of at least 1.024 g mL^-1, 24^) with flow (15 µl min^-1^) for at least 20 hours to induce filamentation and dispersal of the bacteria from the bladder cells. Since the proportion of live filaments is expected to vary with different batches and sources of urine^13^; to avoid variability we used urine from the same batch in experimental replicates. Filaments were collected from the back-opening of the flow chambers and resuspended in LB for fluorescent protein expression, *Live/Dead* cell staining and direct imaging, or stained (FDAA labelling) and fixed.

### Live/Dead cell staining

Staining was performed as recommended by the manufacturer (LIVE/DEAD *BacLight* Bacterial Viability Kit, #L7012, Molecular Probes). Briefly, 3 µl of each dye mixture (SYTO9 and Propidium iodide) was added for each mL sample culture and mixed thoroughly. The dye/sample mixture was incubated for 15 min in the dark at room temperature. After incubation, 2 – 3 µl culture was spotted directly onto agarose pads (1.5% w/v in LB) for imaging. We also confirmed that cells and filaments divided and reverted using only SYTO9 labelling (*Supplementary Figure 3B*).

### FDAA labelling

WT UTI89 filaments were labelled with 1 mM OGDA as described before^35,43^ for 5 minutes and washed twice in PBS, before fixation in 70% ice cool ethanol for 1 hour. After ethanol fixation, cells were washed twice in PBS and placed on agarose pads (1.5% w/v in water) for imaging.

### Microscopy and Imaging

Samples were placed on pre-made agarose (1 - 1.5 % w/w) pads in LB (for dynamics) or M9 (single-molecule imaging) media in 65 µl gene frames (Thermo Scientific, AB0577), left to immobilize and imaged directly.

Live cell epifluorescence and bright field time-lapse imaging of filaments reverting back to rods was done using a Nikon Ti2-E deconvolution microscope with extra-large Field Of View optics (full FOW of the microscope was 25 mm, this ensured us that long filaments could always be imaged within one FOW), equipped with a CFI Plan Apo Lambda DM 100x oil objective (NA 1.45), and an environmental chamber set at 37 °C (Okolab cage incubator). Images were acquired every 2 - 10 minutes as required. Excitation of fluorophores was performed using an Lumencore Spectra II module, and fluorescence was detected using a back illuminated Andor Sona 4.2 sCMOS camera. GFP and mEos3.2 (in the green state) emission was collected through a FITC filter and RFP emission through a Cy5 filter. Filters were from Semrock.

Live cell PALM imaging was performed on a Nikon (Ti2-E) N-STROM v5 with NIS v.5.30 using a 100x 1.49 NA oil objective in TIRF mode. Cover glass slides were washed with 95% EtOH, air dried, cleaned for at least 3 minutes with a plasma cleaner (Harrick plasma, PDC-23G) and used within 15 minutes of cleaning. Imaging was consistently performed at room temperature (∼23 °C). Drift correction during image acquisition was minimised using the integrated PFS4 (Perfect Focus System). 100 nm multi-colour TetraSpeck beads were used as fiducial markers. Prior to single-molecule acquisition, the green state of mEos3.2 was excited by a 488 nm laser to acquire an epifluorescence image using a FITC emission filter cube. For single-molecule acquisition, mEos3.2 was photoconverted to its red state continuously by a 405 nm laser with increasing working powers ranging between 0.1 and 5 W/cm^2^. As readout, mEos3.2 was excited by a 561 nm laser line operating at an average power between 1 and 2 kW/cm^2^. The emission was collected by a quad band (Quad405/488/561/647 filter dual cSTORM). The exposure time was 20 ms and 4000 - 5000 images were typically acquired for each set of images. Images were captured using a sCMOS Flash 4.0 v3 (Hamamatsu) camera.

Note: we also imaged UTI89 cells fixed with 2 % paraformaldehyde using standard protocols^44^, but we noticed that the fixative seemed to interfere with protein localization. This loss of localization was not noticed when imaging live cells, similar to what previous studies with other cell division proteins have found ^45,46^.

### Image analysis

Epi-fluorescence images and movies were visualized and analysed in Fiji^47^. Images and movies were post processed for background subtraction (rolling ball radius 36 pixels). PALM images were processed using the Nikon N-STORM software and Fiji plugin ThunderSTORM^48^ using Gaussian blur of 20 nm for visualization.

### Cell dimensions of rods and filaments during reversal

Cell lengths for rods that had not been through an infection were extracted from bright field images of respective strains. Filaments were followed for one to four divisions on agarose pads using bright field imaging. To simplify the analysis, we only followed cells pinching off from one side from a filament. “Birth” was determined from the first image frame where the poles of cells were not connected. “Lengths at birth” and “symmetry at birth” statistics were generated from length measurements in MicrobeJ^49^ or from manually traced lengths in Fiji. In bright field, a filament was manually classified as dead when it was clearly translucent and had similar grey values as the background. Fluorescence profiles were generated in Fiji and Full Width at Half Maxima (FWHM) were extracted by fitting a gaussian to the generated curves. Plots and statistics were generated using the web apps PlotsOfData^50^ and SuperPlotsOfData^51^, as well as Origin9 pro (Origin Lab, US). Figures were prepared using Adobe Illustrator.

### Western blotting

A volume of cells corresponding to OD_600_ 0.1 was collected from cell cultures. The samples were suspended in loading buffer and resolved by sodium dodecyl sulphate-polyacrylamide gel electrophoresis. Proteins were transferred to nitrocellulose membranes using a semi-dry Transfer-Blot apparatus (Bio-Rad). The membranes were blocked in 5% (w/v) milk and probed with anti-sera to DamX^30^.

## Acknowledgments

The authors would like to thank David Weiss (University of Iowa) for sharing the DamX antisera. BS is funded by a CPDRF scheme from the University of Technology Sydney and the authors acknowledge support via a CAPEX infrastructure grant for a Nikon N-STORM single-molecule microscope. IGD was supported by an Australian Research Council Future Fellowship (FT1601000010).

## Author Contributions

BS and IGD conceptualized the study. DOD contributed reagents and advice on experiments. BS carried out all experiments, and lead the data analysis with input from IGD. BS and IGD wrote the paper with input from DOD.

## Conflict of interest

The authors declare no conflict of interest.

## Data sharing

Data will be available upon request from the authors.

## Notes

### Competing Interest Statement

The authors have declared no competing interest.

## References

1 Stamm, W. E. & Norrby, S. R. Urinary tract infections: disease panorama and challenges. J Infect Dis 183 Suppl 1, S1–4, doi:10.1086/318850 (2001).

2 Wagenlehner, F. M., Tandogdu, Z. & Bjerklund Johansen, T. E. An update on classification and management of urosepsis. Curr Opin Urol 27, 133–137, doi:10.1097/MOU.0000000000000364 (2017).

3 Tandogdu, Z., Cai, T., Koves, B., Wagenlehner, F. & Bjerklund-Johansen, T. E. Urinary Tract Infections in Immunocompromised Patients with Diabetes, Chronic Kidney Disease, and Kidney Transplant. Eur Urol Focus 2, 394–399, doi:10.1016/j.euf.2016.08.006 (2016).

4 Hooton, T. M. Clinical practice. Uncomplicated urinary tract infection. N Engl J Med 366, 1028–1037, doi:10.1056/NEJMcp1104429 (2012).

5 Flores-Mireles, A. L., Walker, J. N., Caparon, M. & Hultgren, S. J. Urinary tract infections: epidemiology, mechanisms of infection and treatment options. Nat Rev Microbiol 13, 269–284, doi:10.1038/nrmicro3432 (2015).

6 Sihra, N., Goodman, A., Zakri, R., Sahai, A. & Malde, S. Nonantibiotic prevention and management of recurrent urinary tract infection. Nat Rev Urol 15, 750–776, doi:10.1038/s41585-018-0106-x (2018).

7 Anderson, G. G. et al. Intracellular bacterial biofilm-like pods in urinary tract infections. Science 301, 105–107, doi:10.1126/science.1084550 (2003).

8 Hannan, T. J., Mysorekar, I. U., Hung, C. S., Isaacson-Schmid, M. L. & Hultgren, S. J. Early severe inflammatory responses to uropathogenic E. coli predispose to chronic and recurrent urinary tract infection. PLoS pathogens 6, e1001042, doi:10.1371/journal.ppat.1001042 (2010).

9 Miller, C. et al. SOS response induction by beta-lactams and bacterial defense against antibiotic lethality. Science 305, 1629–1631, doi:10.1126/science.1101630 (2004).

10 Justice, S. S., Hunstad, D. A., Cegelski, L. & Hultgren, S. J. Morphological plasticity as a bacterial survival strategy. Nat Rev Microbiol 6, 162–168, doi:10.1038/nrmicro1820 (2008).

11 Justice, S. S., Hunstad, D. A., Seed, P. C. & Hultgren, S. J. Filamentation by Escherichia coli subverts innate defenses during urinary tract infection. Proceedings of the National Academy of Sciences of the United States of America 103, 19884–19889, doi:10.1073/pnas.0606329104 (2006).

12 Justice, S. S. et al. Differentiation and developmental pathways of uropathogenic Escherichia coli in urinary tract pathogenesis. Proceedings of the National Academy of Sciences of the United States of America 101, 1333–1338, doi:10.1073/pnas.0308125100 (2004).

13 Andersen, T. E. et al. Escherichia coli uropathogenesis in vitro: invasion, cellular escape, and secondary infection analyzed in a human bladder cell infection model. Infect Immun 80, 1858–1867, doi:10.1128/IAI.06075-11 (2012).

14 McQuillen, R. & Xiao, J. Insights into the Structure, Function, and Dynamics of the Bacterial Cytokinetic FtsZ-Ring. Annu Rev Biophys 49, 309–341, doi:10.1146/annurev-biophys-121219-081703 (2020).

15 Bi, E. F. & Lutkenhaus, J. FtsZ ring structure associated with division in Escherichia coli. Nature 354, 161–164, doi:10.1038/354161a0 (1991).

16 Hale, C. A. & de Boer, P. A. Recruitment of ZipA to the septal ring of Escherichia coli is dependent on FtsZ and independent of FtsA. Journal of bacteriology 181, 167–176 (1999).

17 Addinall, S. G. & Lutkenhaus, J. FtsA is localized to the septum in an FtsZ-dependent manner. Journal of bacteriology 178, 7167–7172 (1996).

18 Hale, C. A. & de Boer, P. A. Direct binding of FtsZ to ZipA, an essential component of the septal ring structure that mediates cell division in E. coli. Cell 88, 175–185 (1997).

19 Haeusser, D. P. & Margolin, W. Splitsville: structural and functional insights into the dynamic bacterial Z ring. Nat Rev Microbiol 14, 305–319, doi:10.1038/nrmicro.2016.26 (2016).

20 Yahashiri, A., Jorgenson, M. A. & Weiss, D. S. Bacterial SPOR domains are recruited to septal peptidoglycan by binding to glycan strands that lack stem peptides. Proceedings of the National Academy of Sciences of the United States of America 112, 11347–11352, doi:10.1073/pnas.1508536112 (2015).

21 Ursinus, A. et al. Murein (peptidoglycan) binding property of the essential cell division protein FtsN from Escherichia coli. Journal of bacteriology 186, 6728–6737, doi:10.1128/JB.186.20.6728-6737.2004 (2004).

22 Arends, S. J. et al. Discovery and characterization of three new Escherichia coli septal ring proteins that contain a SPOR domain: DamX, DedD, and RlpA. Journal of bacteriology 192, 242–255, doi:10.1128/JB.01244-09 (2010).

23 Khandige, S. et al. DamX Controls Reversible Cell Morphology Switching in Uropathogenic Escherichia coli. mBio 7, doi:10.1128/mBio.00642-16 (2016).

24 Iosifidis, G. & Duggin, I. G. Distinct Morphological Fates of Uropathogenic Escherichia coli Intracellular Bacterial Communities: Dependency on Urine Composition and pH. Infect Immun 88, doi:10.1128/IAI.00884-19 (2020).

25 Chen, S. L. et al. Identification of genes subject to positive selection in uropathogenic strains of Escherichia coli: a comparative genomics approach. Proceedings of the National Academy of Sciences of the United States of America 103, 5977–5982, doi:10.1073/pnas.0600938103 (2006).

26 Wegner, A. S., Alexeeva, S., Odijk, T. & Woldringh, C. L. Characterization of Escherichia coli nucleoids released by osmotic shock. J Struct Biol 178, 260–269, doi:10.1016/j.jsb.2012.03.007 (2012).

27 Arends, S. J. R. et al. Discovery and Characterization of Three New <em>Escherichia coli</em> Septal Ring Proteins That Contain a SPOR Domain: DamX, DedD, and RlpA. Journal of bacteriology 192, 242–255, doi:10.1128/jb.01244-09 (2010).

28 Coltharp, C., Buss, J., Plumer, T. M. & Xiao, J. Defining the rate-limiting processes of bacterial cytokinesis. Proceedings of the National Academy of Sciences of the United States of America 113, E1044–1053, doi:10.1073/pnas.1514296113 (2016).

29 Söderström, B. et al. Disassembly of the divisome in Escherichia coli: evidence that FtsZ dissociates before compartmentalization. Molecular microbiology 92, 1–9, doi:10.1111/mmi.12534 (2014).

30 Williams, K. B. et al. Nuclear magnetic resonance solution structure of the peptidoglycan-binding SPOR domain from Escherichia coli DamX: insights into septal localization. Biochemistry 52, 627–639, doi:10.1021/bi301609e (2013).

31 Lyngstadaas, A., Lobner-Olesen, A. & Boye, E. Characterization of three genes in the dam-containing operon of Escherichia coli. Mol Gen Genet 247, 546–554, doi:10.1007/BF00290345 (1995).

32 Tenorio, E. et al. Systematic characterization of Escherichia coli genes/ORFs affecting biofilm formation. FEMS microbiology letters 225, 107–114, doi:10.1016/S0378-1097(03)00507-X (2003).

33 Söderström, B. et al. Coordinated disassembly of the divisome complex in Escherichia coli. Molecular microbiology 101, 425–438, doi:10.1111/mmi.13400 (2016).

34 McCausland, J. W. et al. Treadmilling FtsZ polymers drive the directional movement of sPG-synthesis enzymes via a Brownian ratchet mechanism. Nature communications 12, 609, doi:10.1038/s41467-020-20873-y (2021).

35 Söderström, B., Ruda, A., Widmalm, G. & Daley, D. O. An OregonGreen488-labelled d-amino acid for visualizing peptidoglycan by super-resolution STED nanoscopy. Microbiology (Reading) 166, 1129–1135, doi:10.1099/mic.0.000996 (2020).

36 Kuru, E. et al. In Situ probing of newly synthesized peptidoglycan in live bacteria with fluorescent D-amino acids. Angewandte Chemie 51, 12519–12523, doi:10.1002/anie.201206749 (2012).

37 Mulvey, M. A. et al. Induction and evasion of host defenses by type 1-piliated uropathogenic Escherichia coli. Science 282, 1494–1497, doi:10.1126/science.282.5393.1494 (1998).

38 Wehrens, M. et al. Size Laws and Division Ring Dynamics in Filamentous Escherichia coli cells. Curr Biol 28, 972–979 e975, doi:10.1016/j.cub.2018.02.006 (2018).

39 Söderström, B., Chan, H., Shilling, P. J., Skoglund, U. & Daley, D. O. Spatial separation of FtsZ and FtsN during cell division. Molecular microbiology 107, 387–401, doi:10.1111/mmi.13888 (2018).

40 Lyu, Z. et al. FtsN activates septal cell wall synthesis by forming a processive complex with the septum-specific peptidoglycan synthase in <em>E. coli</em>. bioRxiv, 2021.2008.2023.457437, doi:10.1101/2021.08.23.457437 (2021).

41 Meyer, A. J., Segall-Shapiro, T. H., Glassey, E., Zhang, J. & Voigt, C. A. Escherichia coli “Marionette” strains with 12 highly optimized small-molecule sensors. Nat Chem Biol 15, 196–204, doi:10.1038/s41589-018-0168-3 (2019).

42 Watson, J. F. & Garcia-Nafria, J. In vivo DNA assembly using common laboratory bacteria: A re-emerging tool to simplify molecular cloning. J Biol Chem 294, 15271–15281, doi:10.1074/jbc.REV119.009109 (2019).

43 Hsu, Y. P. et al. Full color palette of fluorescent d-amino acids for in situ labeling of bacterial cell walls. Chem Sci 8, 6313–6321, doi:10.1039/c7sc01800b (2017).

44 Fu, G. et al. In vivo structure of the E. coli FtsZ-ring revealed by photoactivated localization microscopy (PALM). PLoS One 5, e12682, doi:10.1371/journal.pone.0012680 (2010).

45 Buss, J. et al. In vivo organization of the FtsZ-ring by ZapA and ZapB revealed by quantitative super-resolution microscopy. Molecular microbiology 89, 1099–1120, doi:10.1111/mmi.12331 (2013).

46 Buss, J. et al. A multi-layered protein network stabilizes the Escherichia coli FtsZ-ring and modulates constriction dynamics. PLoS genetics 11, e1005128, doi:10.1371/journal.pgen.1005128 (2015).

47 Schindelin, J. et al. Fiji: an open-source platform for biological-image analysis. Nature methods 9, 676–682, doi:10.1038/Nmeth.2019 (2012).

48 Ovesny, M., Krizek, P., Borkovec, J., Svindrych, Z. & Hagen, G. M. ThunderSTORM: a comprehensive ImageJ plug-in for PALM and STORM data analysis and super-resolution imaging. Bioinformatics 30, 2389–2390, doi:10.1093/bioinformatics/btu202 (2014).

49 Ducret, A., Quardokus, E. M. & Brun, Y. V. MicrobeJ, a tool for high throughput bacterial cell detection and quantitative analysis. Nat Microbiol 1, 16077, doi:10.1038/nmicrobiol.2016.77 (2016).

50 Postma, M. & Goedhart, J. PlotsOfData-A web app for visualizing data together with their summaries. PLoS biology 17, e3000202, doi:10.1371/journal.pbio.3000202 (2019).

51 Goedhart, J. SuperPlotsOfData-a web app for the transparent display and quantitative comparison of continuous data from different conditions. Mol Biol Cell 32, 470–474, doi:10.1091/mbc.E20-09-0583 (2021).

