## Supplementary_info_ for "Dynamic localisation of DamX regulates bacterial filamentation and division during UPEC dispersal from host cells"

### Supplementary Figures

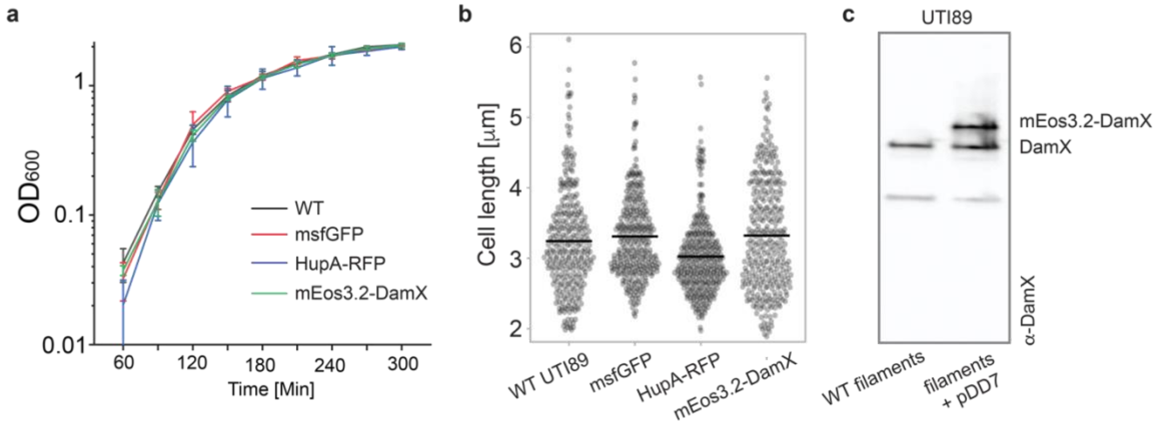


*Supplementary Figure 1. Cell growth and protein detection in UTI89 containing expression plasmids.*

Cell viability measurements of strains and plasmids used in this study: WT UTI89, followed by UTI89 strains expressing msfGFP (pGI5), HupA-RFP (pSTC011) and mEos3.2-DamX (pDD7).

**a**, Growth curves. **b**, Cell lengths for the strains were: WT 3.24 ± 0.75 μm (n = 306), pGI5 3.31 ± 0.6 μm (n = 316), pSTC011 3.02 ± 0.54 μm (n = 340), and pDD7 3.32 ± 0.78 μm (n = 323). **c**, Western Blot detecting DamX protein in dispersed UPEC (including filaments) with and without pDD7. Image quantification indicated that mEos3.2-DamX was produced at 59 ± 0.05 % of total DamX levels with pDD7, calculated using the formula $\left(\frac{Int_{mEos3.2-DamX}}{Int_{mEos3.2-DamX}+Int_{DamX}}\right)$. Measurements are from three biological replicates. Values represent Mean ± SD.

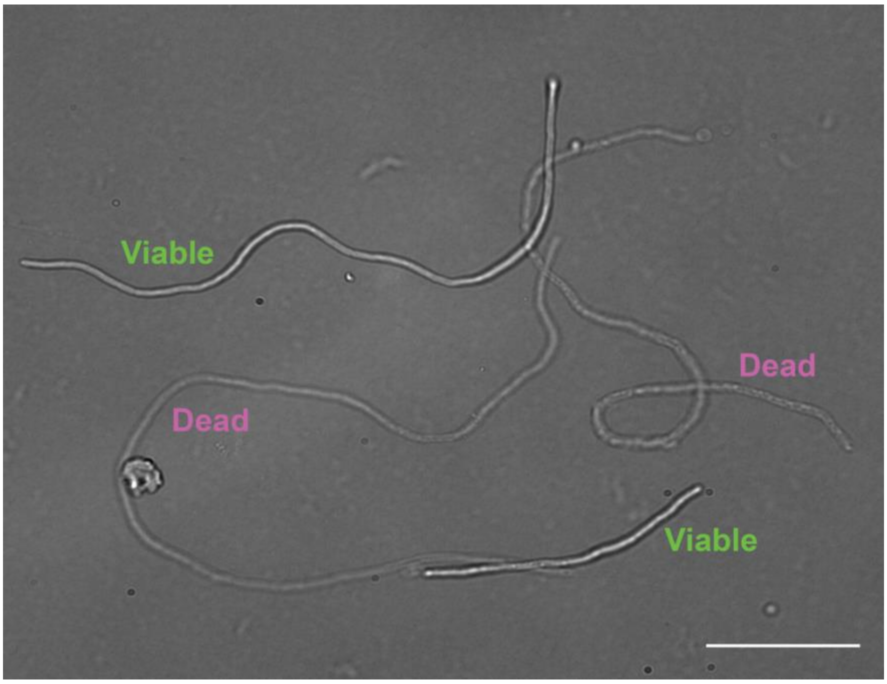


*Supplementary Figure 2. WT UTI89 filaments.*

Bright field image of typical filaments after infection. ‘Viable’ filaments are dense resulting in high contrast while ‘Dead’ filaments are empty membrane shells and therefore highly translucent. Scale bars = 20 $\mu$m.

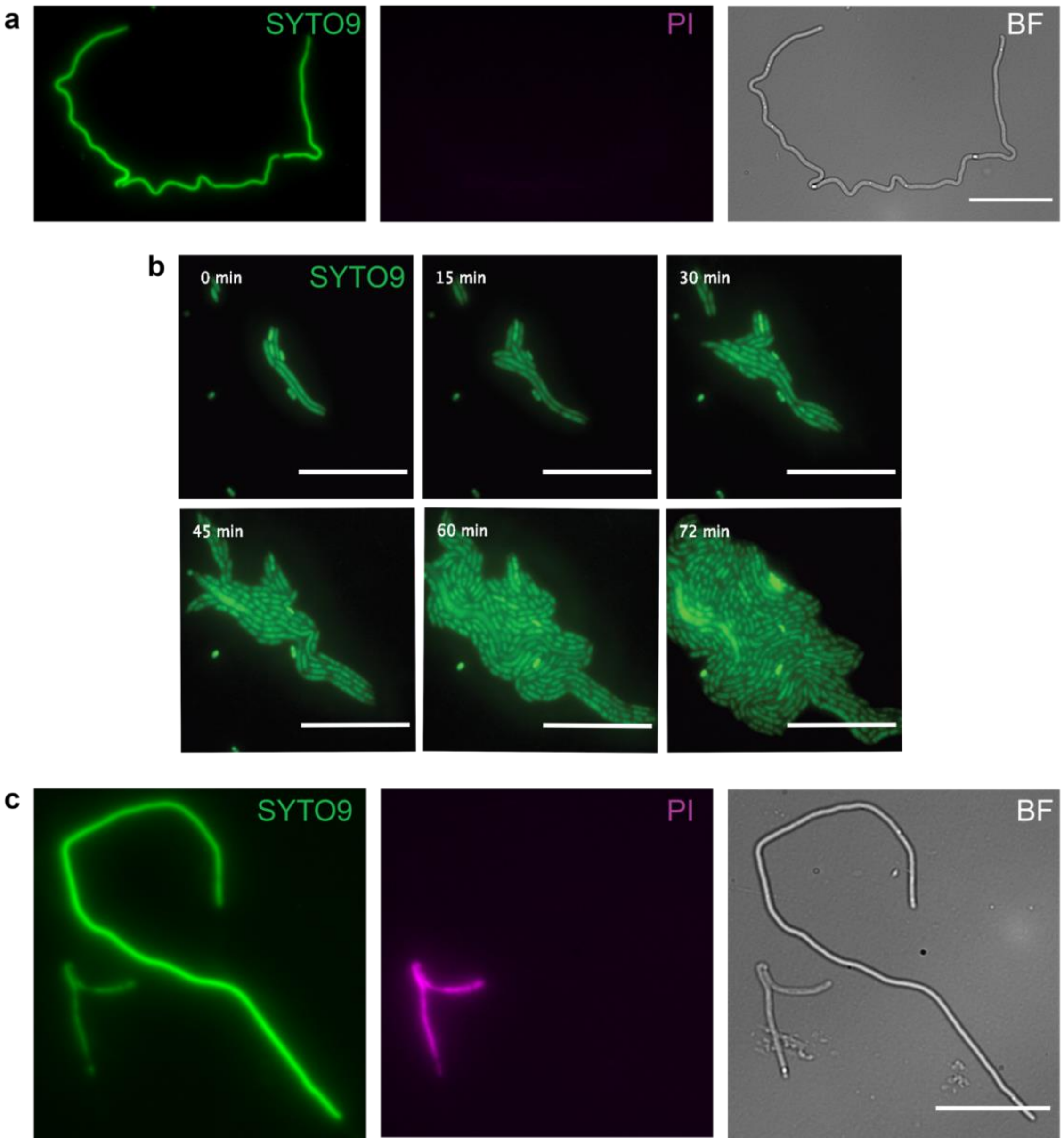


*Supplementary Figure 3. Live/dead staining of UTI89 filaments.*

Filaments were labelled with the *Live/dead Bac*Light staining kit containing SYTO9 and propidium iodide. **a**, Filaments that are alive and show strong contrast in the bright field (BF) image only take up the SYTO9 dye (green). **b**, SYTO9 labelled filamentous and rod cells are viable, and continue dividing rapidly after staining. **c**, Dead filaments show weak contrast in BF and are labelled with propidium iodide (PI), pseudo-coloured magenta. Some dead cells retain some green fluorescence for a time. Scale bars = 20 μm.

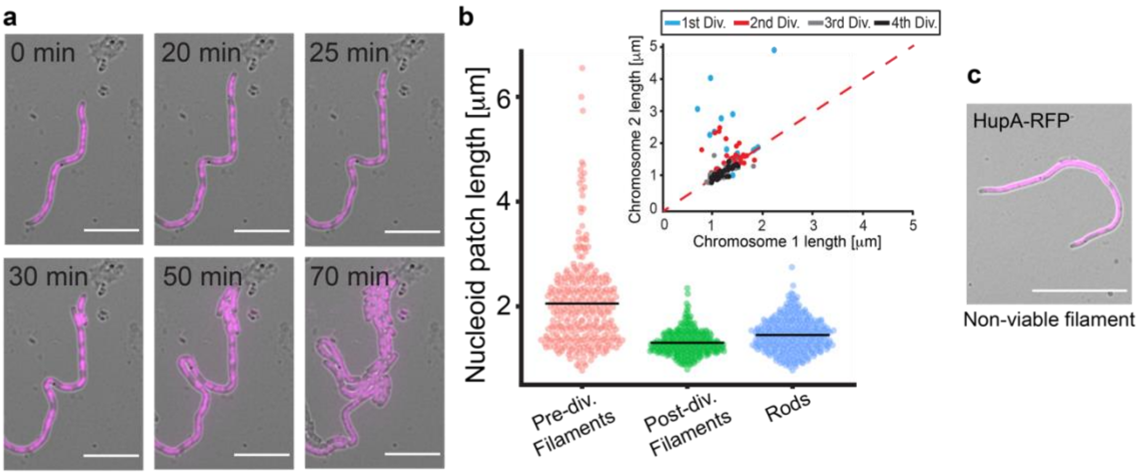


*Supplementary Figure 4. Chromosome organization in filaments reverting to rods.*

**a**, Time-lapse imaging of a representative UTI89 filament expressing HupA-RFP reverting to rods. **b**, Individual nucleoids in live filaments are longer than nucleoids in rod shaped cells. Average chromosome lengths for pre-divisional filaments were 2.05 ± 0.86 µm (red, n = 305), post-divisional filaments were 1.3 ± 0.24 µm (green, n = 254) and for rods that had not gone through an infection 1.45 ± 0.32 µm (blue, n = 308). Values represent Mean ± SD. Inset shows increasing symmetry of daughter chromosomes over divisions (n = 254). Blue dots represent the nucleoid lengths in the cell after the first division of a cell from a filament. Red, grey and black dots represent the nucleoids in subsequent divisions of that rod. The red striped line represents the symmetry line *i.e.*, dots on the line have equally sized chromosomes. Note that chromosome 1 was always picked as the shorter. **c**, A typical non-viable filament with HupA-RFP filling the cytoplasm. Scale bar **a** = 10 µm, **c** = 20 µm.

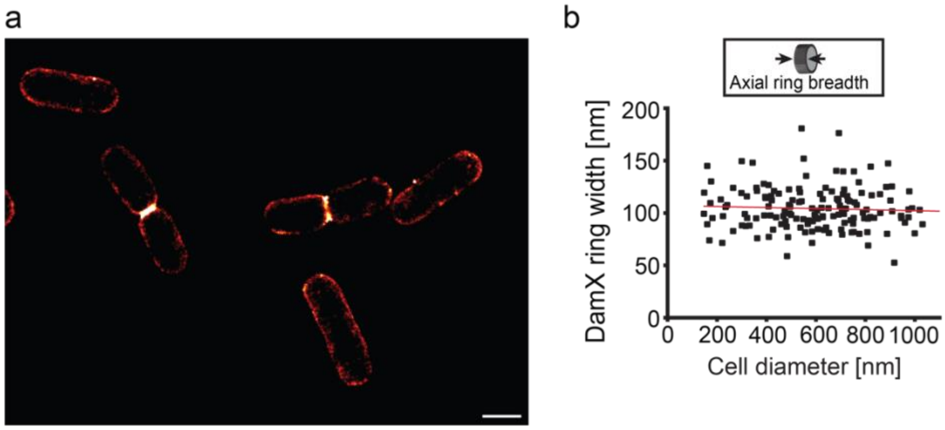


*Supplementary Figure 5. mEos3.2-DamX ring widths in rods*

**a**, Typical PALM images of rod-shaped cells expressing mEos3.2-DamX. Scale bar = 1 μm. **b**, mEos3.2-DamX axial ring breadth during constriction was essentially constant with a mean of 102.5 ± 20.2 nm (n = 150). Values represent Mean ± SD. Red line represents a linear fit to the data, slope of the line was ~ -0.005. Equation of the fitted line: $y = -0.00516\ (\pm 0.00765) * x + 105.4544\ (\pm 4.64415)$

**Supplementary Movie Legends**

Supplementary Movie – SM1.

Upper and lower filaments: Short filaments elongating before reverting back to rods.

Middle filament: initial growth before bursting. Scale bar 10 μm.

Supplementary Movie – SM2.

A long non-viable filament. Scale bar 50 μm.

Supplementary Movie – SM3.

Movie depicts a mixture of filaments; one non-viable long filament and multiple shorter filaments showing initial

elongation before reversal. Scale bar 20 μm.

Supplementary Movie – SM4.

Non-viable filament. Late in the movie are short rods coming from out-of-frame filament reversals observed. Scale bar 20 μm.

Supplementary Movie – SM5.

UTI89 filaments labelled with Live/Dead stain. Green indicates live cells, magenta indicates dead cells. Scale bar 20 μm.

Supplementary Movie – SM6.

Reversal of an UTI89 filament after a round of infection. Scale bar 10 μm.

Supplementary Movie – SM7.

Division of rod-shaped cells expressing mEos3.2-DamX from two different infection samples. Scale bar 4 μm.

Supplementary Movie – SM8.

*De novo* assembly of mEos3.2-DamX in a filament during reversal. Scale bar 10 μm.

Supplementary Movie – SM9.

*De novo* assembly of mEos3.2-DamX in a filament during reversal. Scale bar 10 μm.
